# Lineage transcription factors co-regulate subtype-specific genes providing a roadmap for systematic identification of small cell lung cancer vulnerabilities

**DOI:** 10.1101/2020.08.13.249029

**Authors:** Karine Pozo, Rahul K. Kollipara, Demetra P. Kelenis, Kathia E. Rodarte, Xiaoyang Zhang, John D. Minna, Jane E. Johnson

**Affiliations:** Department of Neuroscience, UT Southwestern Medical Center, Dallas, TX 75390, USA; Department of Surgery, UT Southwestern Medical Center, Dallas, TX 75390, USA; McDermott Center for Human Growth and Development, UT Southwestern Medical Center, Dallas, TX 75390, USA; Department of Oncological Sciences, Huntsman Cancer Institute, University of Utah, Salt Lake City, UT; Hamon Center for Therapeutic Oncology Research, UT Southwestern Medical Center, Dallas, TX 75390, USA; Simmons Comprehensive Cancer Center, UT Southwestern Medical Center, Dallas, TX 75390, USA; Department of Internal Medicine, UT Southwestern Medical Center, Dallas, TX 75390, USA; Department of Pharmacology, UT Southwestern Medical Center, Dallas, TX 75390, USA

## Abstract

Lineage-defining transcription factors (LTFs) play key roles in tumor cell growth, making them highly attractive, but currently “undruggable”, small cell lung cancer (SCLC) vulnerabilities. Delineating LTF genomic binding sites and associated chromatin features would provide important insights into SCLC dependencies. Here we map super-enhancers (SEs) across multiple patient-derived SCLC preclinical models, and find SE patterns are sufficient to classify the models into the recently defined, LTF-based, SCLC subtypes. 3D-chromatin conformation analysis identified genes associated with SEs that define subtype-specific tumor signatures with genes functioning in diverse processes. Focusing on ASCL1-high SCLC (SCLC-A), we found ASCL1 physically interacts with NKX2-1 and PROX1. These factors bind overlapping genomic regions, and co-regulate a set of genes, including genes encoding cell surface proteins, SCN3A and KCNB2 enriched in SCLC-A. Genetic depletion of NKX2-1 or PROX1 alone, or in combinations with ASCL1, did not inhibit SCLC growth more than that achieved by depleting ASCL1 alone. We demonstrate the SE signature supports the LTF classification of SCLC, identify NKX2-1 and PROX1 as ASCL1 co-factors, and substantiate the central importance of ASCL1 as a key dependency factor in the majority of SCLC. The LTF and SE gene sets provide a molecular roadmap for future ASCL1 therapeutic targeting studies.

## INTRODUCTION

Small cell lung cancer (SCLC) is an aggressive and incurable form of lung cancer labeled as a recalcitrant cancer by the NCI and US Congress (1). Patients exhibit dramatic responses to initial platin doublet chemotherapy, but in nearly all cases, SCLCs become resistant to this treatment with tumor recurrence (2). Despite extensive genomic analyses of SCLC tumors, which identified the near ubiquitous inactivation of tumor suppressors TP53 and RB1, and co-occuring high expression of a MYC family member (MYCL, MYCN, MYC), no effective therapeutic vulnerabilities (such as mutant EGFR found in some lung adenocarcinomas) have been identified (1, 3, 4). Inter-tumor heterogeneity has been identified for SCLC and expression of lineage-defining transcription factors (LTFs, such as ASCL1, NEUROD1, POU2F2, and YAP1) appear to play a major role in this heterogeneity (5). The majority of SCLC express the LTFs ASCL1 and/or NEUROD1, and also express a panel of neuroendocrine genes (1). In addition, ASCL1 and NEUROD1 drive distinct oncogenic programs (and are referred to as lineage oncogenes), thereby conferring different molecular and physiological properties to ASCL1- and NEUROD1-containing SCLC (1, 6). Other subsets of SCLC do not express neuroendocrine makers (and are referred to as non-neuroendocrine) and are dependent on other LTFs, POU2F3 (7) or YAP1. These findings led to a current classification where expression of different LTFs define four molecular SCLC subtypes, i.e., SCLC-A, SCLC-N, SCLC-P and SCLC-Y, where A, N, P, and Y stand for ASCL1, NEUROD1, POU2F3 and YAP1, respectively (5). Separating SCLC patients’ tumors into molecular subtypes may provide therapeutic information, and a better understanding of the mechanism(s) the LTFs play in the pathogenesis of SCLC is needed to improve diagnosis and treatment.

The ASCL1-expressing SCLCs, SCLC-A, is the predominant SCLC subtype (1, 4). ASCL1 is a basic helix-loop-helix TF and belongs to a network of TFs that regulate the expression of genes supporting SCLC growth and survival (6). During development, ASCL1 is essential for the differentiation of pulmonary neuroendocrine cells (PNECs) (8-10), a cell lineage from which most SCLC are thought to originate (11, 12). Inactivation of TP53 and RB1 in PNECs or their progenitors appear to be initiating events for SCLC (13, 14). PNECs are rare neuroendocrine cells and are found either in isolation or as small clusters of 20-30 cells at bronchial branch points (15). They represent only ∼1% of the lung epithelium, and in diseases such as asthma, and in smokers, PNEC numbers can increase (10). PNECs function in oxygen sensing, modulation of immune response, and the repair of damaged lung epithelium upon airway damage (16, 17). A small subpopulation of PNECs can divide to repair the damaged epithelium via a Notch-dependent mechanism and return to quiescence following RB1/TP53 activation, and dysfunctions in this process may underlie SCLC tumorigenesis (16, 18). Consistently, ASCL1 is expressed in PNECs and in SCLC tumors where it commands the expression of neuroendocrine lineage genes such as *CHGA, CALCB, INSM1* and genes underlying SCLC progression such as *RET*, and NOTCH-signaling pathway genes (6). Genetic deletion of *Ascl1* in mouse models precludes PNEC formation (8, 9), and importantly, deletion of *Ascl1* in a genetically engineered mouse model (GEMM) of SCLC prevents SCLC tumorigenesis (6). Thus, ASCL1 is a potential therapeutic target for SCLC. Despite its central importance in SCLC tumorigenesis, mechanisms controlling ASCL1 transcriptional activity in SCLC, and the downstream oncogenic program that ASCL1 controls need to be fully characterized to provide therapeutically relevant information.

Deregulation of transcriptional activity and downstream gene expression underlies cancer formation and development (19). Core lineage-driving TFs such as ASCL1 regulate the expression of lineage-specific genes, in part by cooperating with co-factors, and are often found enriched at transcriptionally active genomic regions called super-enhancers (SE). These regions contain abundant TF binding sites, transcriptional molecular machinery, and can be identified using the epigenetic mark H3K27 acetylation (H3K27ac) (20, 21). In cancer cells, SEs are associated with oncogene expression (20, 22-25), and targeting SE components, such as BRD4 and CDK7, inhibits cancer progression (26-28). Thus, identifying SEs across SCLC subtypes, provides key information on cooperating TF networks and their tumor-specific gene expression programs (29). SE information also helps identify subtype-specific vulnerabilities as demonstrated in medulloblastoma (30). Indeed, ASCL1, NEUROD1 and POU2F3, and their respective downstream targets, are associated with SE in some patient derived SCLC lines (6, 7, 28). Here we use a comprehensive approach to study SCLC SEs and identify subtype-specific features of SCLC with the twin goals of delving deeper into SCLC biology, and working out a roadmap for SCLC therapeutic targeting (31, 32).

To address how ASCL1 and other TFs cooperate to support SCLC survival and growth we focused on the SCLC-A subtype, investigating how ASCL1 cooperates with other TFs to drive SCLC. We identify a physically interacting network of TFs that includes ASCL1, NKX2-1 and PROX1, and show that they co-regulate common transcriptional targets in SCLC-A that regulate Notch-signaling and neuroendocrine identity, with NKX2-1 also enriched at cell-cycle genes. In addition, the cell-surface protein encoding genes *SCN3A* and *KCNB2* are downstream targets of ASCL1, NKX2-1 and PROX1, and provide potential SCLC-A biomarkers.

## RESULTS

### Super Enhancer (SE) identification in SCLC cell lines and PDXs support SCLC molecular subtype classifications SCLC-A, SCLC-N, and SCLC-P

Tumor-promoting genes are associated with H3K27ac-rich genomic regions computationally defined as SEs (20, 21), and lineage oncogenes ASCL1 and NEUROD1 are found associated with SEs in some patient-derived SCLC lines (6, 28). To investigate whether SEs across many preclinical models of SCLC would support the recently proposed molecular classification (5), we interrogated a panel of SCLC lines and patient-derived xenografts (PDXs) representing the 3 most common SCLC subtypes in this classification, SCLC-A, SCLC-N, and SCLC-P (Fig. 1A) with H3K27ac chromatin immunoprecipitation followed by sequencing (ChIP-seq). SEs were defined computationally (21), and unbiased clustering of these SEs, and SEs from other studies including 4 non-small cell lung carcinoma (NSCLC) lines and one human broncho-epithelial cell line (HBEC), revealed that the human SCLC preclinical models (with one exception) distributed into 3 distinct SCLC SE clusters (Fig. 1B,C). These SE clusters correlate with ASCL1, NEUROD1 and POU2F3 expression providing further definition of the SCLC-A, SCLC-N and SCLC-P subtypes.

**Figure 1.**
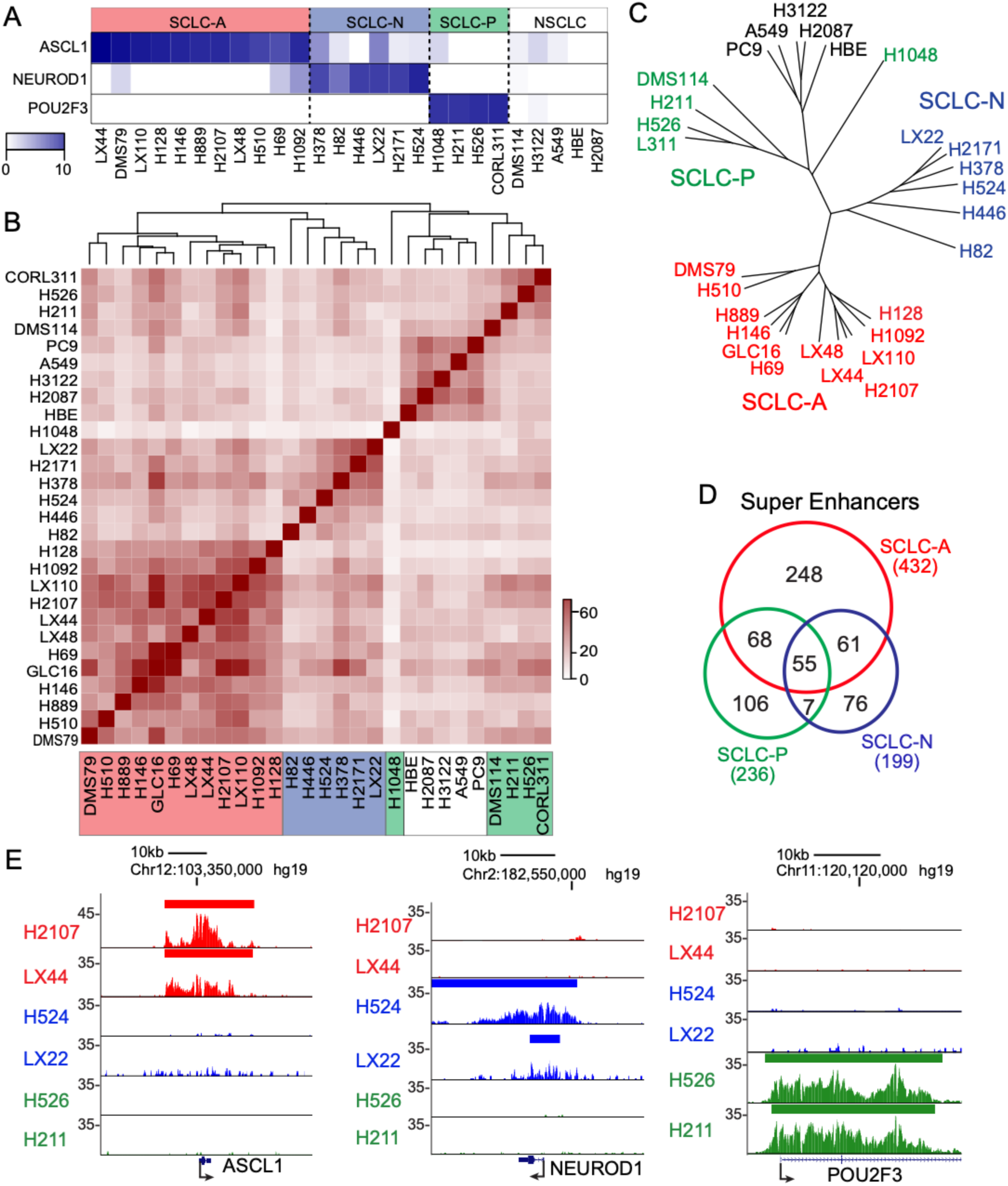
Identification of SEs in SCLC cell lines and PDXs support SCLC molecular subtype classifications SCLC-A, SCLC-N, and SCLC-P. (A) Heat-map from RNA-seq showing ASCL1, NEUROD1 and POU2F3 gene expression in the SCLC cell lines and PDXs used in this study. (B-C) Unbiased cluster analysis (B, heat map) and (C, dendrogram) shows that except for NCI-H1048, the SEs can be used to classify SCLC cell lines and PDXs into the same subtypes recently proposed (5). (D) VENN diagram representing the number of SEs unique to or shared between SCLC subtypes. For a SE to be included in the analysis, it had to be present in 6 of the 11 SCLC-A, 4 of the 6 SCLC-N, or 3 of the 5 SCLC-P samples. (E) Representative genome tracks showing the H3K27ac ChIP-seq signal around *ASCL1, NEUROD1*, and *POU2F3*. The bar above the track indicates regions computationally defined as SEs. Data from SCLC cell lines are color-coded; SCLC-A (red), SCLC-N (blue), SCLC-P (green). LX22, LX44, LX48, and LX110 are PDX. All other samples are cell lines. Also see Supplemental Table S1 for SE genomic coordinates.

To identify SEs characteristic of each SCLC TF subtype, we included SEs found in the majority of samples from any given model (such as 6 of 11 SCLC-A models, 4 of 6 SCLC-N models, or 3 of 5 SCLC-P models), resulting in 432 (ASCL1), 199 (NEUROD1), and 236 (POU2F3) SEs, respectively (Fig. 1D). Within the SCLC-A subtype, 248 SE are unique while 61 SE are shared with the SCLC-N subtype, and 68 with the SCLC-P subtype. A total of 55 SEs are shared between the three subtypes consistent with some common gene expression features of these related tumors (Fig. 1D). Thus, the LTFs, in addition to their characteristic high expression defining their subtype, are also associated with SEs found only in that subtype (Fig. 1E). Furthermore, for *ASCL1* and *NEUROD1*, which are small, 2 exon genes, the H3K27ac enrichment that identified the SE is across the entire gene body. These results show that SE-defined subgroups are also consistent with the current use of the LTFs to classify SCLC (5), and that these factors are indicators of distinct chromatin landscapes.

### 3D chromatin conformation using HiChiP identifies SCLC-A AND SCLC-N subtype SE-associated genes many of which are associated with NE processes

SE-associated genes are enriched in cell identity and disease relevant genes, and thus, are candidates for playing important functions in SCLC pathogenesis. To identify SE-associated genes in the SCLC-A and the SCLC-N subtypes, we combined the H3K27ac ChIP-seq data used to identify SE with data from a 3D chromatin conformation assay, HiChIP, a protein-centric chromatin conformation method, that identifies genes whose transcription start sites are found in proximity to the SE (33). The SEs represent transcriptionally active chromatin regions while the HiChIP identified chromatin loops that connect these SEs to their target genes. Using the work-flow shown in Fig. 2A, that incorporates RNA-seq expression data from SCLC preclinical models, we identified SE-associated genes for SCLC-A and SCLC-N (Supplemental Table S1). (Note: Because we do not have HiChIP data for SCLC-P, the SCLC-P SE-associated genes are those predicted using GREAT algorithms that largely represent SE neighboring genes (34). In addition, in all cases we used a gene expression filter based on RNA-seq from the relevant SCLC preclinical models. The SE-associated genes identified include ASCL1, NEUROD1, and POU2F3, as well as other TFs, many of which have been associated with important processes in SCLC such as INSM1, NFIB, FOXA2 and MYC (4, 6, 7, 28, 31, 35-41) (Fig. 2B).

**Figure 2.**
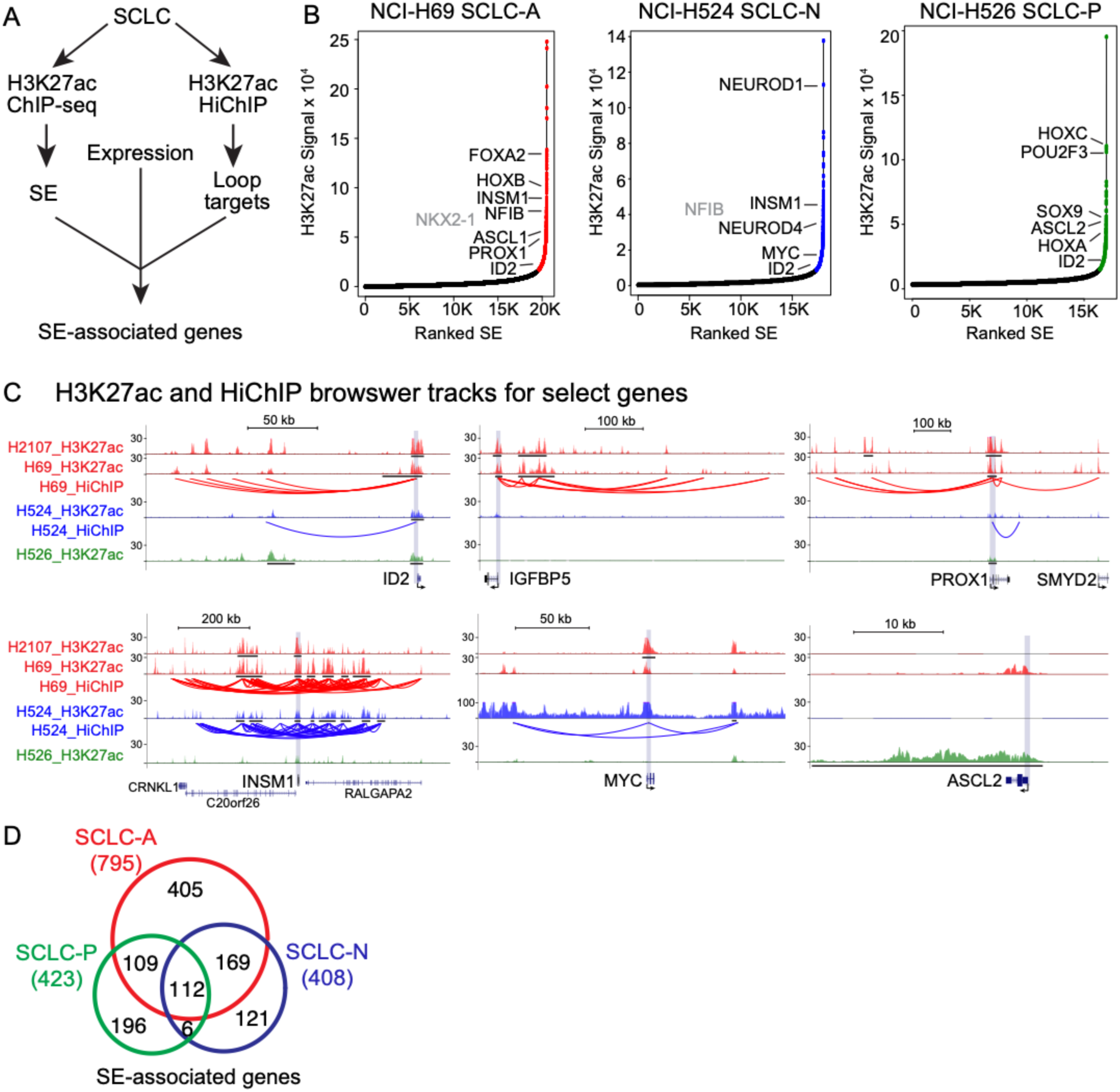
Identification of SE-associated genes in SCLC subtypes. (A) The work-flow used to identify expressed SE-associated genes including H3K27ac ChIP-seq, HiChIP, and RNA-seq. (B) Visualization of the H3K27ac ChIP-seq signal in a representative SCLC cell line from each subtype showing those that meet the SE criteria. Some TF encoding genes associated with the SEs are highlighted for comparison. *NKX2-1* and *NFIB* are in gray because in the particular cell line shown, they just missed the stringent criteria used. (C) Human genome tracks showing the H3K27ac signal and HiChIP data associated with representative SE-associated genes shared and distinct across the cell lines. The black bars under the tracks indicate a region identified as an SE. (D) Venn diagram showing the number of SE-associated genes that are unique to or shared between the 3 SCLC subtypes. Total number of SE-associated genes is indicated. For an SE to be included in the analysis, it had to be present in 6 of the 11 SCLC-A, 4 of the 6 SCLC-N, or 3 of the 5 SCLC-P samples. An expression cut off of 5 FPKM in at least one relevant cell line was used as an expression filter. Data from SCLC cell lines are color-coded; SCLC-A (red), SCLC-N (blue), SCLC-P (green). See Supplemental Table S1.

Analyses of the identified SE-associated genes for the different SCLC subtypes revealed many distinct and shared features across the SCLC subtypes and comprise genes involved in diverse biological processes. SE-associated genes (997 genes total) meeting our criteria were detected including 795 in SCLC-A, 408 in SCLC-N, and 423 in SCLC-P. Among these genes, 405 (51%), 121 (30%) and 196 (46%) were unique to the SCLC-A, -N, -P subtypes respectively, while 112 (14-27%) were common to the 3 subtypes (Fig. 2D, Supplemental Table S1). The genome browser tracks show regulatory elements defined by the H3K27ac ChIP-seq signal as well as chromatin loops identified from the HiChIP data. These data highlight genes uniquely identified such as IGFBP5 in SCLC-A, MYC in SCLC-N, and ASCL2 in SCLC-P, and those shared such as INSM1 in SCLC-A and SCLC-N, or ID2 in all 3 subtypes (Fig. 2C). Overall the 3D chromatin conformation HiChIP provide even greater detail on differences and commonalities between lineage TF SCLC subtypes and provide a resource for systematic determination of which of the products of these genes are potential therapeutic vulnerabilities for each SCLC subtype.

### PROX1 and NKX2.1 are identified in a transcriptional complex with ASCL1, physically interact, and share binding profiles in the genomes of SCLC-A cell lines

There is substantial evidence supporting a key requirement for ASCL1 in promoting the survival and growth of SCLC (6, 8, 42, 43). The *ASCL1* gene is not only associated with SEs in SCLC, but ASCL1 protein itself is also enriched in these genomic regions (6). Transcriptional complexes are often required to add DNA binding specificity and optimal transcriptional activity, thus we needed to determine what proteins are part of an ASCL1 transcriptional complex. To understand how ASCL1 function is regulated in SE regions, we set out to identify other TFs that may be found physically together with ASCL1. ASCL1 was immunopurified from SCLC-A subtype NCI-H2107 cell nuclear extracts and the immune pellet was analyzed by mass-spectrometry. It is known that ASCL1 heterodimerizes with class I bHLH factors called E-proteins, encoded by the genes *TCF3, TCF4*, and *TCF12*, for efficient DNA binding and transcriptional activity (44). Valildation of our approach came when we identified the known ASCL1 E-protein heterodimeric partners TCF3 (previously E12/E47), TCF4 (previously E2-2), and TCF12 (previously HEB), in these complexes, as well as detecting 271 other proteins (Fig. 3A, Supplemental Table S2). Of these 271 proteins, 53 are encoded by SE-associated genes in SCLC-A cells including transcription related factors such as NKX2-1, PROX1, TRIM28, TCF12, CBX3, CBX5, and SMARCC2. We chose to focus on two homeodomain-containing TFs, NKX2-1 (also known as TTF1) and PROX1, because they have regulatory roles in epithelial lung cell biology, and thus may be particularly relevant to SCLC-A pathogenesis. NKX2-1 is highly expressed in thyroid cancer and SCLC (45), while PROX1 plays an oncogenic role in different cancers including SCLC (46). To validate the mass spectrometry-identified ASCL1/NKX2-1 and ASCL1/PROX1 interactions, ASCL1 was immunoprecipitated from NCI-H2107 cells and surveyed by immunoblotting (Fig. 3B). NKX2-1 and PROX1 were both detected in the immune pellet, although NKX2-1 was less abundant than PROX1. In further validation of ASCL1 specific interactions, we detected neither factor in the SCLC-N cell line NCI-H524. Surveying RNA expression of *NKX2-1* and *PROX1* across SCLC tumors (4) and SCLC lines (43) we found both factors are more consistently expressed in SCLC than in NSCLC. In addition, *NKX2-1* expression was enriched in SCLC-A subtypes while *PROX1* was less specific for the SCLC-A subtype (Fig. 3C). These results indicate that NKX2-1 and PROX1 interact in a complex with ASCL1, and raise the obvious hypothesis that ASCL1, PROX1 and NKX2-1 might act together to regulate the pathogenesis of SCLC-A.

**Figure 3.**
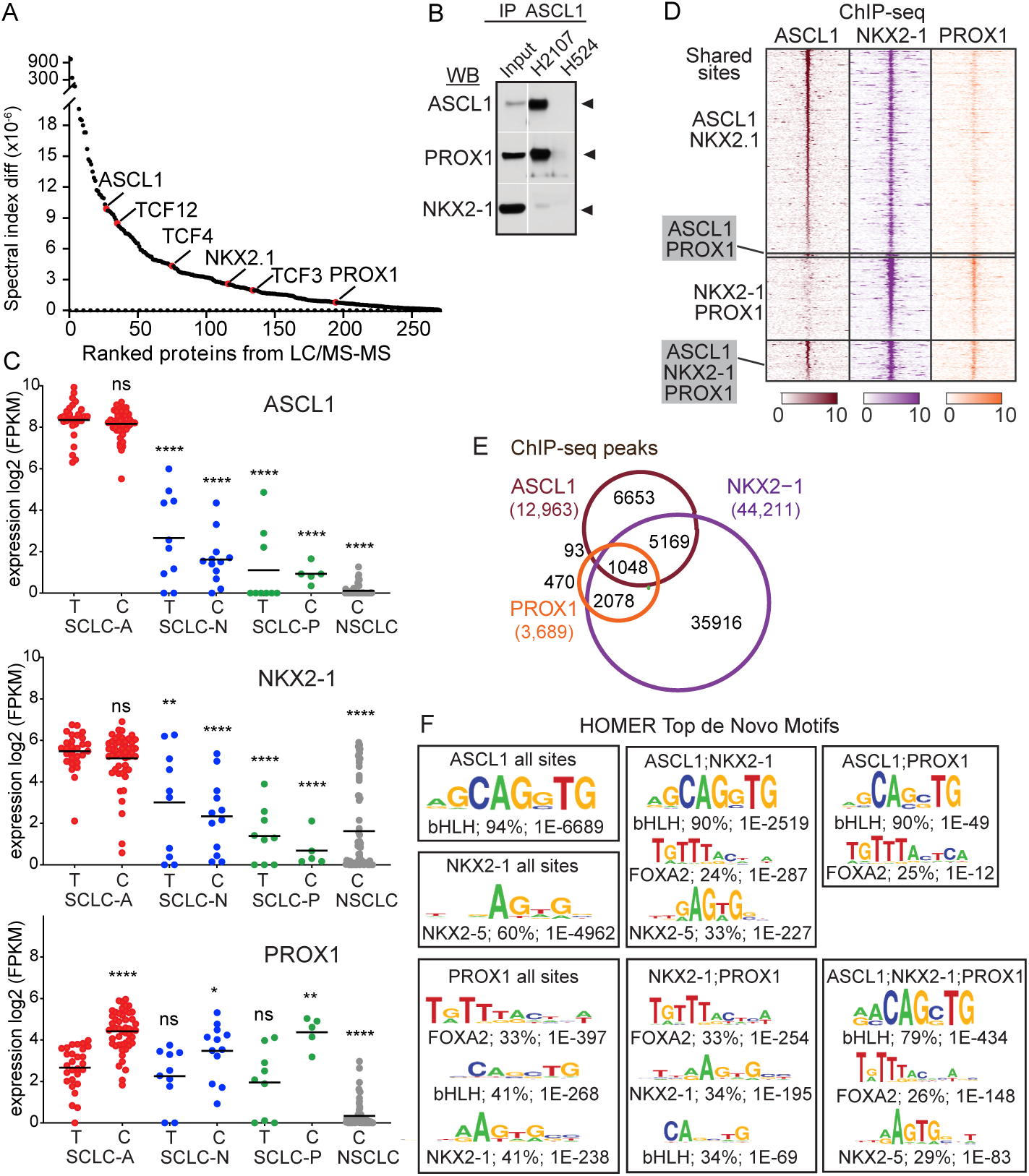
PROX1 and NKX2-1 co-immunoprecipitate with ASCL1 and have overlapping binding profiles in the genomes of SCLC-A cell lines. (A) Proteins identified by mass spectroscopy of ASCL1 immunoprecipitates from NCI-H2107 cells shown ranked by spectral index. ASCL1 and its E-protein cofactors TCF3, TCF4, and TCF12 are shown, as are NKX2-1 and PROX1. See Supplemental Table S2 for a complete list of proteins identified. (B) Immunoblots showing PROX1 and NKX2-1 co-immunoprecipitate with ASCL1 from NCI-H2107 cells. The SCLC-N cell line NCI-H524 does not express ASCL1 and was used as a negative control. Input is the nuclear fraction from NCI-H2107. None of the TFs were detected in the NCI-H524 nuclear fraction (not shown). (C) RNA-seq (log2 FPKM) showing expression of *ASCL1, NKX2-1*, and *PROX1*, in SCLC tumors (T) or cell lines (C) in the specified SCLC subtype or NSCLC. The line indicates the mean and the asterisks indicate p-values for each sample compared to SCLC-A tumors. * p<0.05, ** p<0.01, **** p<0.0001. (D) ChIP-seq heat map and (E) VENN diagram for ASCL1, PROX1 and NKX2.1 peaks identified in at least 2 of 3 SCLC-A cell lines (NCI-H2107, NCI-H128, and NCI-H889) identifies unique and shared genomic binding sites. See Supplemental Table S3 for peak coordinates and Supplemental Table S4 for TF-binding associated genes. (F) Top *de novo* motifs identified by HOMER in ChIP-seq peak regions. The text indicates the motif enriched, the factor family predicted to bind the motif, the percent incidence of the motif in ChIP-seq peak regions, and the p-value significance of the specified motif.

To determine whether ASCL1, PROX1 and NKX2-1 regulate a common set of gene targets in SCLC-A, we mapped their binding sites in the genomes of 3 SCLC-A cell lines by ChIP-seq. Overall, 1,048 genomic sites were bound by all three factors. In addition, many binding sites were identified for each TF including 12,963 for ASCL1, 3,689 for PROX1, and 44,211 for NKX2-1 (Fig. 3D, E) (Supplemental Table S3, S4). The co-enrichment for binding motifs for the 3 TFs also supports substantial overlap in genomic regions bound. HOMER was used to identify the top *de novo* motifs within the ChIP-seq peaks (47). The motifs known for ASCL1 and NKX2-1 were found enriched in their respective ChIP-seq data with 94% of ASCL1 ChIP-seq peaks containing the bHLH motif, and 60% of NKX2-1 ChIP-seq peaks containing a motif most similar to the related factor NKX2-5 (Fig. 3F), consistent with these factors directly binding specific DNA sequences. In contrast, PROX1 ChIP-seq peaks were similarly enriched for a FOXA2 motif (33%), a bHLH motif (41%), and an NKX2-1 motif (41%), a finding more consistent with PROX1 being recruited as a cofactor to these sites. In ChIP-seq bound regions shared by all three factors, or ASCL1 and either of the other two factors, the most enriched *de novo* binding motif is the bHLH (79-90%) (Fig. 3F), and the NKX2-1/5 and FOXA2 motifs are found within 24-34% of these shared sites. The presence of the FOXA2 motif suggests that FOXA2 might form a DNA-protein complex with ASCL1, PROX1 and/or NKX2-1, however, FOXA2 did not co-immunoprecipitate with these TFs in lysates from NCI-H2107 cells (data not shown). Overall the *de novo* motif analysis is consistent with the existence of ASCL1/PROX1, ASCL1/NKX2-1 and ASCL1/NKX2-1/PROX1 containing DNA-protein complexes in SCLC-A cells.

### Cross-regulatory relationships between ASCL1, PROX1, and NKX2-1

ASCL1, PROX1, and NKX2-1 are associated with SEs in SCLC-A cells (Fig. 2B). In addition, we discovered through ChIP-seq that each factor binds to regions surrounding its own gene and to the genes encoding the other factors (Fig. 4A), setting up the possibility for auto- and cross-regulatory relationships between these TFs in SCLC-A. To investigate functional dependencies between ASCL1, PROX1 and NKX2-1, each TF was knocked-down individually or in combination in NCI-H2107 cells using siRNAs (Fig. 4B). Immunoblots probing protein levels of each factor were used to assess the functional dependencies between factors. We found that knock down of NKX2-1 led to decreases in ASCL1 and PROX1, knock down of ASCL1 led to decreased PROX1, and knock down of PROX1 led to a modest decrease in ASCL1 (Fig. 4B,B’,D). Thus, these findings support potential cross-regulatory relationships between the factors, but the robustness of the cross regulation is not predictable from the ChIP-seq data (Fig. 4A). To determine the effect of a decrease in the TF individually or in combination, we also assessed cell viability after siRNA. A decrease of ASCL1 reduced SCLC viability by ∼50%, whereas knock down of NKX2-1 or PROX1 had more modest effects (20-25%) (Fig. 4C). Importantly, combined knock down of two or more of these TFs did not enhance the decrease in NCI-H2107 cell viability over ASCL1 knock down alone.

**Figure 4.**
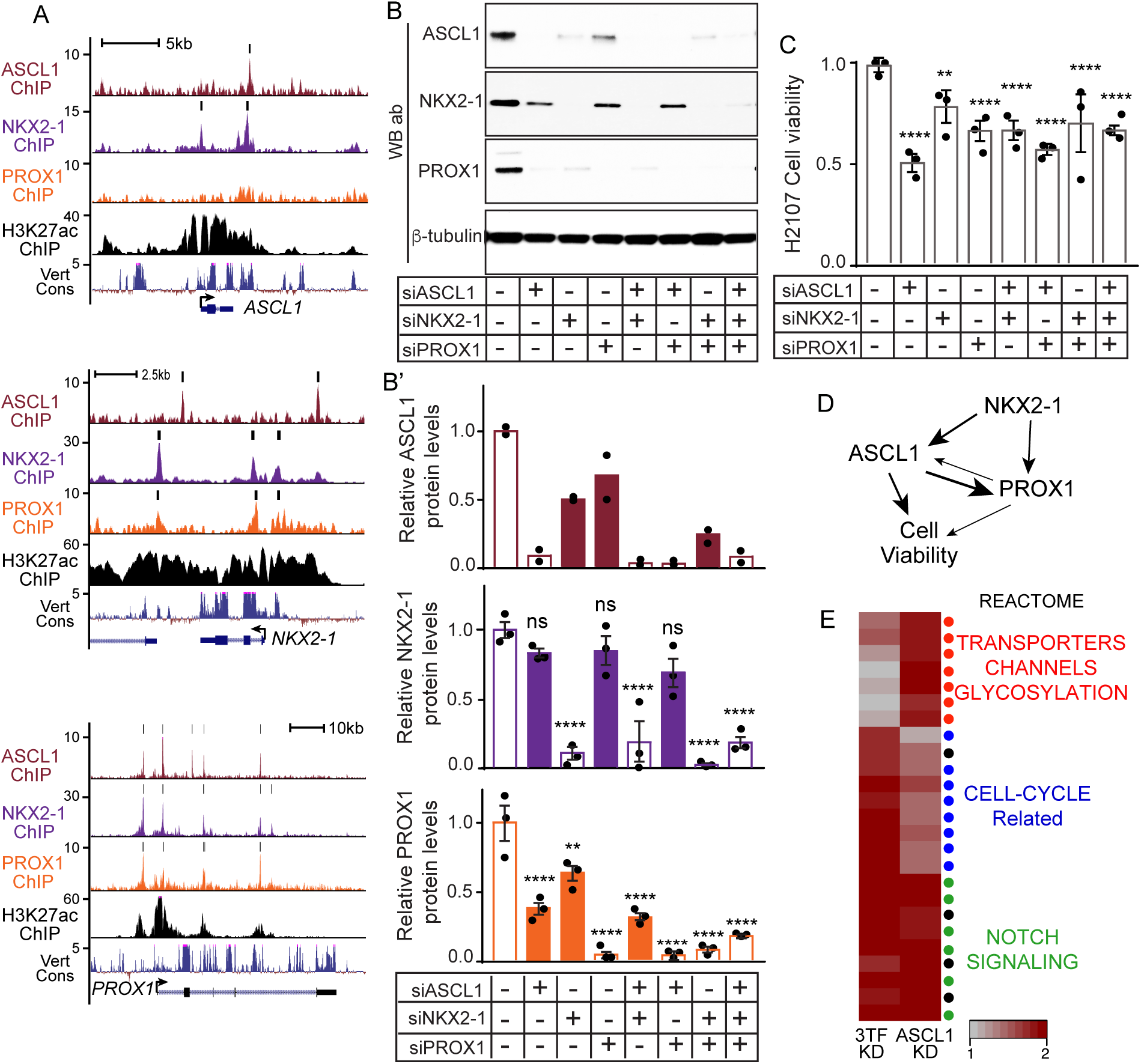
Cross-regulatory relationships between ASCL1, PROX1, and NKX2-1. (A) UCSC Genome Browser tracks showing ASCL1, NKX2-1, PROX1 and H3K27ac ChIP-seq signals in NCI-H2107 cells around the genes encoding *ASCL1, NKX2*-1, and *PROX1*. Black bars above the tracks indicate computationally called binding sites. (B) Immunoblots and (B’) quantification to assess protein levels, and (C) WST-1 assay for cell viability after siRNA-mediated knock down of ASCL1, PROX1, NKX2-1 individually or in combination in NCI-H2107 cells harvested at 72 hours post-transfection. Each data point represents a biological replicate, error bars indicate SEM. ANOVA with Bonferroni’s multiple comparisons test was used to determine significant differences relative to WT, p-values (**p<0.01, ****p<0.0001, ns=not significant). (D) Diagram summarizing functional dependencies between the three factors. (E) Enrichment plot showing normalized enrichment scores for significantly enriched gene sets (FDR <= 0.05) in down regulated genes upon ASCL1 and 3TF knock down in NCI-H2107 as in (B), compared to controls. Heat map illustrates genes activated by ASCL1 are enriched in gene sets for NOTCH signaling (green) and cell-cycle (blue) but also include transporters and ion channels (red). The 3 TF knock down further alters cell-cycle related genes. See Supplemental Table S5 for the detailed list of DEG genes and pathway analysis.

### Genes directly regulated by ASCL1, PROX1, and NKX2-1 in SCLC

To discover genes regulated by ASCL1 versus those co-regulated by the three TFs together, we performed RNA-seq from the knock down experiments in NCI-H2107 cells with control siRNA, siASCL1, and the combination siASCL1;siNKX2-1;siPROX1 (n=2 for each) assayed at 72 hours after transfection. Differential gene expression (DEG) was determined relative to control siRNA samples (Fig. 4E, Supplemental Table S5). Consistent with known functions of ASCL1 in regulating NOTCH pathway genes, the DEGs reflecting activation by ASCL1 were those enriched in NOTCH related pathways. These were not changed with combinatorial depletion of the co-factors NKX2-1 and PROX1. Genes encoding transporters, ion channels, and glycosylation-related proteins are also regulated by ASCL1 but these effects were modulated by the additional loss of NKX2-1 and PROX1. Notably, although ASCL1 activates cell-cycle related genes, NKX2-1 and PROX1 enhance this activity resulting in a robust decrease in multiple cell-cycle related genes when the 3 TFs were knocked down in combination. The TF repressed genes were not enriched in any specific pathway. (See Supplemental Table S5 for the detailed list of DEG genes and pathway analysis).

Taken together, the protein-protein binding and the substantial overlap in genomic binding sites for ASCL1, NKX2-1, and PROX1, suggest they have overlapping functions and some cross-regulatory relationships. Disrupting ASCL1 levels alone has a profound effect on NOTCH pathway genes while the combined decrease of ASCL1, NKX2-1, and PROX1 additionally alters the transcriptome by disrupting expression of a cohort of cell-cycle related genes.

### Identifying downstream targets of the ASCL1/NKX2-1/PROX1 TF network

The overlap in binding of ASCL1, NKX2-1, and PROX1 TFs in the SCLC-A genome suggests a common set of gene targets that contribute to the growth or other characteristics of these cancer cells. To identify the unique and common direct transcriptional targets of these factors, we identified the genes associated with the TF ChIP-seq peaks using the 3D chromatin conformation from SCLC-A HiChIP data. We defined 3,253 ASCL1-associated targets where 95% are shared with NKX2-1-associated targets. A third of these, 1,011, are also shared with PROX1-associated targets (Fig. 5A,B and Supplemental Table S4). We further refined the list of target genes using functional dependencies based on differential gene expression in response to siRNA knock down of ASCL1 alone or in combination with NKX2-1 and PROX1 in NCI-H2107 cells. Focusing first on ASCL1 downstream targets, there were 1,340 DEGs with 751 identified as being activated by ASCL1 and 589 genes repressed (Fig. 5C, Supplemental Table S5). Of these, 295 were associated with an ASCL1 ChIP-seq binding site indicating direct regulation, and 64% of these are activated by ASCL1, consistent with ASCL1’s known activity as a transcriptional activator. The most highly enriched gene ontology terms describing the ASCL1 targets are those associated with catecholamine biosynthesis, NOTCH pathway, and nervous system development (Fig. 5E, Supplemental Table S6). These findings support the known relationship between ASCL1 activity and modulation of NOTCH signaling, the balance of which controls cell proliferation and lineage-differentiation in neural and neuroendocrine stem/progenitor cell niches (18, 48-51), and ASCL1-mediated regulation of neural and neuroendocrine lineage genes, particularly in development (6, 52-55).

**Figure 5.**
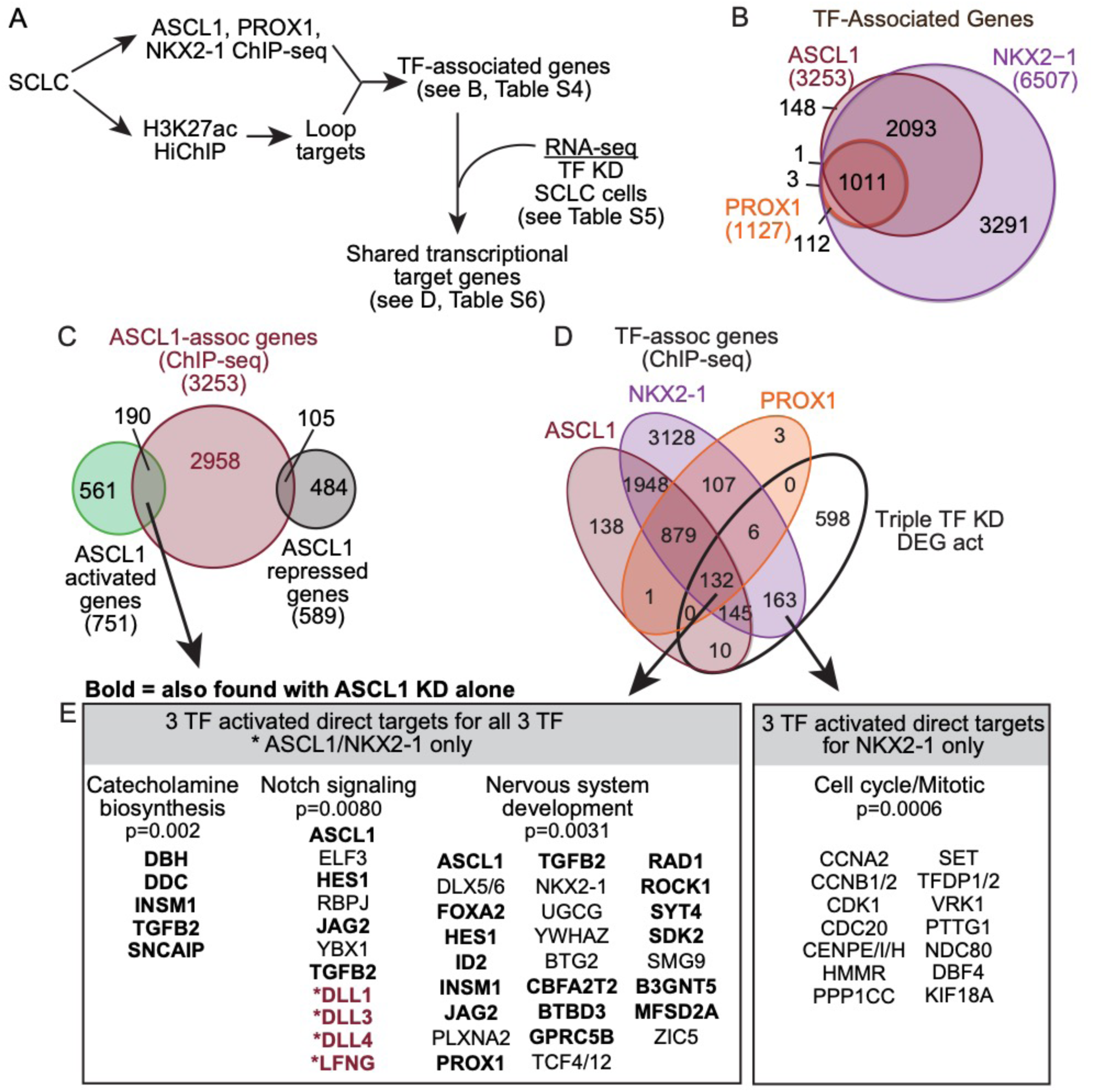
Transcriptional targets of ASCL1, NKX2.1 and PROX1. (A) The work-flow used to identify transcriptional targets of the three TFs using ChIP-seq, HiChIP, and RNA-seq. (B) VENN showing the extensive overlap in TF-associated genes identified from the ChIP-seq and HiChIP analysis. (C) VENN showing the subset of ASCL1 ChIP-seq associated genes (from B) that are differentially expressed (DEG) in NCI-H2107 cells with siASCL1 knock down followed by RNA-seq. (D) VENN from (B) but with the DEG from NCI-H2107 cells with siASCL1;siNKX2-1;siPROX1 knock down followed by RNA-seq. Data is shown for TF activated genes only. (E) GO pathway analysis of genes activated by the 3 TFs show an enrichment for terms associated with NOTCH signaling for ASCL1, and cell cycle related genes for NKX2-1. See Supplemental Table S6 for the full lists of direct targets of ASCL1 and all 3TFs, and the pathway analysis.

We next asked whether the combined loss of all three TFs, ASCL1, NKX2-1, and PROX1, would enhance the ASCL1 knock down phenotype or not, and whether any additional pathways might be affected. Here we compared the DEGs identified in the triple TF knock down, with all the TF-associated genes identified through ChIP-seq. From this analysis there were two groups of genes that were enriched with gene ontology terms (Fig. 5D,E, Supplemental Table S6). First, 132 genes were associated with binding of all three TFs, with interactions supported by HiChIP data, and were significantly decreased in the triple knock down. These strongly overlapped with the genes associated with catecholamine biosynthesis, NOTCH signaling, and nervous system development as was seen with ASCL1 alone (Fig. 5E bolded genes were identified with ASCL1 alone). The second notable finding is that many cell cycle/mitosis related genes appear to be directly activated by NKX2-1 since they are associated with NKX2-1 ChIP-seq peaks but not those from ASCL1 or PROX1 (Fig. 5D,E). From these analyses, we conclude that ASCL1 is key to regulating NOTCH signaling and neuroendocrine features in SCLC, while NKX2-1 and PROX1 as co-factors may largely function to increase the robustness of this activity. NKX2-1 appears to have additional, specific activity in regulating some cell-cycle genes independent of ASCL1 and PROX1.

### The ion channels, SCN3A and KCNB2, are downstream targets of the NKX2.1/ASCL1/PROX1 transcriptional network providing new SCLC-A cell surface biomarkers

The 132 genes we identified in SCLC-A cells that are both associated with SEs and regulated by the core TF network (differentially regulated with genetic knockdown) provide a source list of targets or biomarkers for this subtype of SCLC (Supplemental Table S6). Two examples are genes encoding the ion channels, SCN3A and KCNB2, which were identified as direct targets of ASCL1, NKX2-1, and PROX1, had high signals for H3K27ac in NCI-H2107, and *SCN3A* mRNA was significantly enriched in SCLC-A models (Fig. 6A,B). To test if *SCN3A* and *KCNB2* are dependent on ASCL1, NKX2.1 and PROX1 for expression, their protein levels were assessed by immunoblot in NCI-H2107 cell lysates after siRNA-mediated knock down of each of the TFs, individually. Knock down of each factor alone resulted in a dramatic decrease in SCN3A and KCNB2 protein relative to treatment with a control siRNA (Fig. 6C), demonstrating the requirement for all three of these TF for *SCN3A* and *KCNB2* expression in SCLC-A cells. We tested whether the loss of these ion channels contributed to the decreased SCLC viability seen with *ASCL1* knock down. Efficient knock down of *SCN3A* or *KCNB2* in NCI-H2107 did not significantly decrease cell viability by WST-1 assay (Fig. 6D-F). However, pharmacological inhibition of KCNB2 by Quinine resulted in a modest decrease in colony formation in soft agar with NCI-H889 cells (SCLC-A), compared with no effect detected with NCI-H524 (SCLC-N) or NCI-H526 (SCLC-P) cells (Fig. 6G). Inhibition of SCN3A with the blocker ICA-121431 did not significantly affect colony formation in SCLC-A, -N, or -P cell lines (data not shown). These findings demonstrate expression of these two ion channels is dependent on ASCL1, NKX2-1, and PROX1, however, they are largely dispensable for SCLC survival and growth in the assays used here.

**Figure 6.**
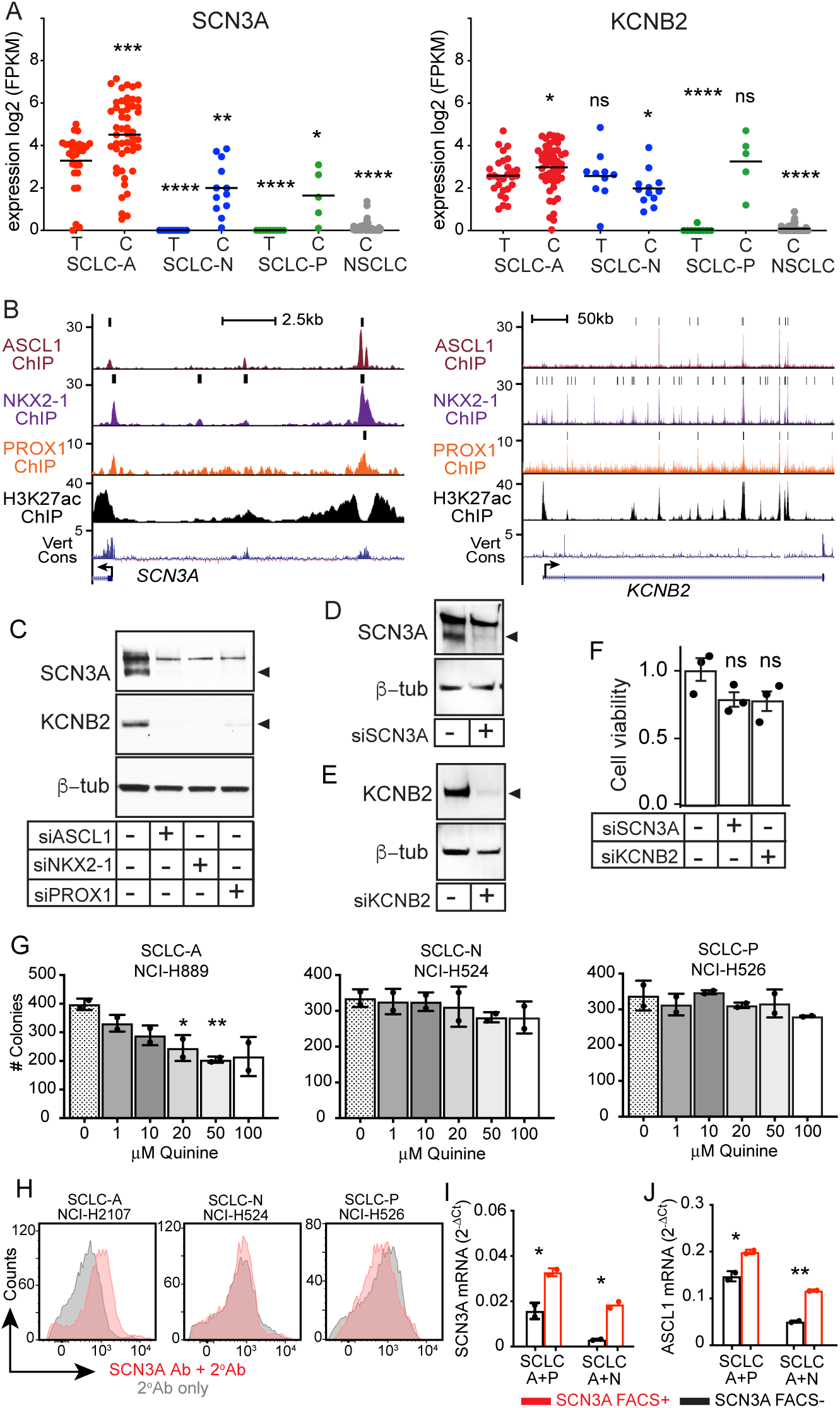
Common transcriptional targets of ASCL1, NKX2.1 and PROX1 include the ion channels *SCN3A* and *KCNB2*. (A) RNA-seq (log2 FPKM) showing expression of *SCN3A* and *KCNB2*, in SCLC tumors (T) or cell lines (C) in the specified SCLC subtype or NSCLC. The line indicates the mean, and the asterisks indicate p-values for each sample compared to SCLC-A tumors. * p<0.05, ** p<0.01, *** p<0.001, **** p<0.0001. (B) UCSC Genome Browser tracks showing ASCL1, NKX2-1, PROX1 and H3K27ac ChIP-seq signals in NCI-H2107 cells around the genes encoding *SCN3A* and *KCNB2*. Black bars above the tracks indicate computationally called binding sites. (C-E) Immunoblots showing SCN3A and KCNB2 protein in NCI-H2107 cells, 72 hours post-transfection with control siRNAs or siRNA targeted against either *ASCL1, NKX2-1*, or *PROX1* (C), or *SCN3A* (D) or *KCNB2* (E). (F) WST-1 assay for cell viability from cells from (D,E). Each data point represents a biological replicate, and error bars indicate SEM. ANOVA with Bonferroni’s multiple comparisons test was used to determine significant differences relative to control. ns; not significant. (G) Quantification of colony formation assays in soft agar using varying doses of KCNB2 inhibitor Quinine shows increased sensitivity of SCLC-A NCI-H889, compared to SCLC-N NCI-H524, and SCLC-P NCI-H526. (H) Histograms showing SCN3A+ cells are detected in live SCLC-A but not SCLC-N or SCLC-P cell cultures using an SCN3A extracellular domain specific antibody (red). The background fluorescence with the secondary antibody only is shown (gray). (I,J) RT-qPCR for *SCN3A* (I) and *ASCL1* (J) mRNA from FACs isolated SCN3A+ (red) or SCN3A-(black) cells from mixtures of SCLC-A with SCLC-P or – N. Each data point represents a biological sample, error bars = SD around mean, unpaired t-test, * p<0.05, ** p<0.01.

We then tested whether, as a cell-surface expressed ion channel, SCN3A could be valuable as a biomarker for identification or manipulation of the SCLC-A subtype. Using a commercially available antibody raised to the extracellular domain of SCN3A, FACs analyses identified live SCN3A+ cells in SCLC-A but not SCLC-N or SCLC-P cell cultures (Fig. 6H). Furthermore, FACS with this SCN3A antibody was able to effectively isolate live SCN3A+ cells from mixtures of SCLC-A + SCLC-N or SCLC-A + SCLC-P cells (Fig. 6I,J). Thus, SCN3A as a biomarker can be used for purifying SCLC-A cells that will be of value in studies of SCLC plasticity (e.g. transition of SCLC from one LTF expressing phenotype to another) and also may provide value for direct targeting of antibody-based therapeutic loads to these SCLC-A cells in a tumor.

## DISCUSSION

A better understanding of molecular subtypes and their underlying biology is necessary to develop new treatments for SCLC. Several approaches ranging from gene expression analysis to systems biology have been used to classify SCLC tumors (4) (1) (32) (56) and a consensus classification based on lineage TF expression of ASCL1, NEUROD1, POU2F3, and YAP1 has been established (5). To get a more comprehensive understanding of SCLC lineage TF phenotypes and to establish a roadmap of potential therapeutic targets, we profiled whole genome SEs in a number of patient derived SCLC cell and xenograft preclinical models. SEs are large genomic regions enriched in TF-binding sites that regulate the expression of genes for cell fate and identity in normal cells. Cancer cells can acquire additional SEs by genomic rearrangements (23, 24, 57-59). Genes associated with SEs are expressed at higher levels and thought to promote carcinogenesis (20, 26), and as a result, SE components are being evaluated as potential therapeutic targets in many cancers including SCLC (28). Unbiased clustering of preclinical SCLC and NSCLC models according to their SEs, distributed the SCLC models into 3 groups that were characterized by the expression of one of the lineage-defining TFs, ASCL1, NEUROD1 and POU2F3 and thus corresponding to the SCLC-A, SCLC-N and SCLC-P subgroups defined in the consensus classification (5). Because of their small numbers, we did not analyze the SCLC-Y subgroup in this study. Noteworthy, SE analyses showed that SCLC PDXs and cell lines clustered similarly, indicating SCLC lines conserve typical SCLC molecular features despite being maintained in cell culture. Further integrated analyses that were focused on the SCLC-A phenotype revealed enrichment of SE-target genes related to Notch signaling, nervous system development and catecholamine biosynthesis, consistent with a neuroendocrine phenotype and a previous study showing a link between the neuroendocrine phenotype and the Notch pathway (4). Overall our study substantiates that SEs can be used to classify tumors, a strategy used to classify other tumor types such as medulloblastoma (30), and demonstrates the ASCL1, NEUROD1, POU2F3, YAP1 consensus classification, which is based on the predominant expression of a single lineage transcription factor, also represents distinct chromatin landscapes.

SEs inform on the core TF regulatory circuitry and in particular help identify interconnected TF networks (19). Here, by integrating SE profiling with proteomics and TF-ChIP-seq, we found that the TFs ASCL1, NKX2-1 and PROX1 are associated with SEs in the SCLC-A subtype, have physical and functional dependencies, and share common transcriptional targets, and as such define a TF regulatory network. Our findings provide experimental support for the TF network and downstream targets predicted from gene expression correlations extracted from SCLC_CellMiner (https://discover.nci.nih.gov/SclcCellMinerCDB) (Tlemsani et al., 2020).

NKX2-1 is a master regulator TF involved in pulmonary epithelial cell differentiation (60). NKX2-1 is also found in most SCLCs (61) where its SCLC expression correlates with that of ASCL1 (32, 40). Our findings that ASCL1 and NKX2-1 interact and bind overlapping sites on the genome in SCLC are consistent with Hokari et al. (2020). They found that ASCL1 and NKX2-1 exogenously expressed in human kidney HEK293 cells, immunoprecipitate and bind to adjacent sites (62). In the current study, we provide data showing this interaction also occurs endogenously in SCLC. Together our studies, with those of Hokari et al., support ASCL1/NKX2-1 cooperation in SCLC-A. Likewise, there are reports of PROX1 involvement in neurodevelopment and cancer (46, 63). Here we detected PROX1 in SCLC models and found that it complexes with ASCL1 in SCLC, and decreasing PROX1 decreases SCLC viability.

ASCL1, NKX2-1 and PROX1 share overlapping binding sites on the genome and common downstream targets. Gene ontology analysis identifies genes enriched in Notch signaling and catecholamine biosynthesis. This highlights the strong association of ASCL1 with Notch signaling found throughout neural development and in NE lineages (18, 51, 52, 64). It also highlights the function of this TF network in NE lineage identity. NKX2-1 together with ASCL1 is involved in the reprogramming of normal epithelial cells into NE SCLC (65). Furthermore, PROX1 has previously been found to regulate the expression of secretory-granule related genes, including chromatogranin A (66), which are stored and released with catecholamines in/from dense core vesicles. These findings are consistent with ASCL1, NKX2-1 and PROX1 cooperating to regulate the NE phenotype of SCLC.

Horie et al. have proposed that ASCL1 and NKX2-1 regulate the expression of NFIB, a gene important for the progression of SCLC to a metastatic state (37, 67, 68). This conclusion was based on findings that NKX2-1 and NFIB expression is correlated in SCLC, and that NFIB levels are reduced upon NKX2.1 knockdown. We find NFIB is a downstream transcriptional target of ASCL1, PROX1, and NKX2-1 by ChIP-seq and RNA-seq (Tables S4, S5). However, NFIB is still expressed in NKX2-1-negative SCLCs, and NFIB protein levels are not changed upon knockdown of either ASCL1 or NKX2-1. Thus, ASCL1 regulation of NFIB seems to be context dependent.

Expression interdependencies by siRNA-mediated TF knock down suggests an NKX2-1/ASCL1 axis (Fig. 4D). We found NKX2-1 modulates ASCL1 levels but ASCL1 is not required for NKX2-1 expression. This is in contrast to Horie et al. (2018) who made the opposite observation, while Hokari et al. (2020) found that knockdown of either ASCL1 or NKX2-1 caused the other to increase. While these discrepancies need to be resolved by additional studies, multiple experiments support some cross-regulatory relationship between ASCL1 and NKX2-1. The direction of this cross-regulation is uncertain and might depend on the SCLC cell/tumor or other intrinsic molecular factors.

By integrating SE profiling with proteomics and other functional genomics in SCLC-A cells, we identified gene products potentially enriched in the SCLC-A subtype. For the subsequent validation steps, we focused on targets encoding cell surface proteins because the latter are attractive targets for antibody-based targeted therapies, e.g. (69), cell targeted therapy (70), and potential markers for molecular subtyping of SCLC tumors from liquid biopsy for patient stratification (71). We selected two genes encoding ion channels, *SCN3A* and *KCNB2*, to interrogate further. SCN3A is a voltage-gated sodium channel found in neuroendocrine cancers (72), and is reported to be involved in proliferation and metastasis (73). KCNB2 is a voltage-gated potassium channel detected in gastric cancer cell lines (74). We found enrichment of these ion channels, particularly *SCN3A*, in cell lines and tumors from the SCLC-A subtype. While, knock-down of either KCNB2 or SCN3A had no effect on viability of SCLCs in vitro, and only pharmacological inhibition of KCNB2 showed any specific inhibition in a colony formation assay, these two proteins may be clinically useful as biomarkers, or as targets for antibody-based targeted therapies or cell targeting antigens.

In conclusion, we developed a multilayered, integrative approach to investigate SEs related to ASCL1, NEUROD1, and POU2F3, and probe the transcriptional mechanisms underlying the SCLC-A subtype. We provide a valuable list of SCLC SE subtype-specific genes to exploit in the future for molecular diagnostic or therapeutic applications for precision medicine approaches.

## METHODS

### Cell lines

The cell lines used in this study were NCI-H2107, NCI-H889, NCI-H510, NCI-H69, NCI-H1092, NCI-H378, NCI-H524, NCI-H446, NCI-H2171, NCI-H82, NCI-H1048, NCI-H211, and NCI-H526 (Hamon Cancer Center Collections, UT Southwestern Medical Center, Dallas, TX). Cell lines were authenticated by DNA-fingerprinting using the PowerPlex 1.2 Kit (Promega DC6500), and tested for mycoplasma using an e-Myco Kit (Boca Scientific 25235). Cells were maintained in RPMI-1640 media (ThermoFisher Scientific 11875135) supplemented with 5% heat deactivated fetal bovine serum (Sigma Aldrich) without antibiotics at 37°C in a humidified atmosphere containing 5% CO2 and 95% air.

### Patient-derived xenografts (PDX) model establishment

PDX models JHU-LX22, JHU-LX-44, JHU-LX-48 and JHU-LX-110 were established by using dissociated cells from resected PDX tumors (gift from C. Rudin and J.T. Poirier, Memorial Sloan Kettering Cancer Center, NYC) (75). Cells (1×10^6^) were centrifuged, resuspended 1:1 (vol/vol) in matrigel and HITES media (Corning) and injected subcutaneously in the flank of NGS mice. Animals were sacrificed 6-8 weeks after implantation. Tumors were collected and processed for ChIP as described by (76).

### siRNA transfections and cell survival assays

For transfections, NCI-H2107 cells were plated at a density of 3×10^5^ cells/ml in 12-well plates and transfected by adding dropwise 25-100 nM siRNA mixed to 20 μl X-Treme siRNA transfection reagent (Sigma-Aldrich 4476093001) in Opti-MEM™ reduced serum medium (ThermoFisher Scientific) as per manufacturer’s recommendations. The siRNAs used for this study were NKX2-1 MISSION® siRNA (Sigma-Aldrich HS0100189848), PROX1 siRNA (Santa Cruz Biotechnology sc-106451), ASCL1 siRNA (ThermoFisher Scientific 36758255), SCN3A siRNA (Sigma-Aldrich HS0200316523), KCNB2 siRNA (Sigma-Aldrich HS0200338274), control siRNA-A (Santa Cruz Biotechnology sc-37007) and MISSION® siRNA Universal Negative Control #1 (Sigma-Aldrich SIC001). Cell viability was measured 72 hours post-transfection using the WST-1 cell proliferation reagent (Sigma-Aldrich5015944001). Cells were resuspended and transferred to a 96-well plate (90 μl), supplemented with 10 µl cell proliferation reagent and incubated at 37°C in a humidified atmosphere containing 5% CO2 and 95% air for 2 hours. Absorbance was read using a multi-well plate reader. Each experiment was done in triplicate.

### RNA extraction for sequencing

Cells were harvested by centrifugation and pellets were resuspended in Trizol® (ThermoFisher Scientific 15596026). Total RNAs were extracted using Arcturus PicoPure RNA (Thermofisher Scientific KIT0204) isolation kit as per the manufacturer’s recommendation. RNA were checked using a Bioanalyzer and sequenced on a NextSeq 550 DX Illumina sequencing platform

### Chromatin immunoprecipitation (ChIP) and libraries preparation

H3K27ac ChIPs were performed as described by (76). Briefly, ∼7 ×10^7^ cells were fixed in 1% formaldehyde (EMS 50980485) for 10 minutes at room temperature. Nuclear extracts were prepared by resuspending cell pellets in Lysis Buffer 1 (50 mM HEPES-KOH, pH 7.5, 140 mM NaCl, 1 mM EDTA, 10% glycerol, 0.5% Igepal, 0.25% Triton X-100) supplemented with protease inhibitors, followed by centrifugation and resuspension in Lysis Buffer 2 buffer (10 mM Tris-HCl, pH 8.0, 200 mM NaCl, 1 mM EDTA, 0.5 mM EGTA) supplemented with protease inhibitors. After centrifugation, pellets were resuspended in Lysis Buffer 3 (10 mM Tris-HCl, pH 8.0, 100 mM NaCl, 1 mM EDTA, 0.5 mM EGTA, 0.1% Na-Deoxycholate, 0.5% N-lauroylsarcosine) supplemented with protease inhibitors and sonicated in a Diagenode Bioruptor for 40 minutes, 30sec:30sec on:off cycles, high power. Chromatin (200 μg) was incubated overnight with 10 μg rabbit anti-H3K27ac (Abcam Ab4729) in ChIP buffer (20 mM Tris-HCl, pH 8.0, 150 mM NaCl, 1 mM EDTA, 0.1% Triton X-100) supplemented with protease inhibitors. This step was followed by incubation with 35 μl protein A/G agarose plus (50% slurry) (ThermoFisher Scientific 20421) for 6 hours. After 6 washes in RIPA buffer (50 mM HEPES-KOH, pH 7.6, 500 mM LiCl, 1 mM EDTA, 1.0% NP-40, 0.7% Na-Deoxycholate) followed by 1 wash in TEN buffer (10 mM Tris-HCl, pH 8.0, 1 mM EDTA, 50 mM NaCl), immunoprecipitated DNA was eluted in buffer E (50 mM Tris-HCl, pH 8.0, 10 mM EDTA, 1.0% SDS) for 30 min at 70°C and reverse cross-linked for 12 hours at 65°C. DNA was treated with proteinase K and RNAse A and DNA purified using QIAgen PCR purification kit. Transcription factor ChIPs were conducted as described above except that eluted DNA was treated with proteinase K prior to reversed cross-link for 15 hours at 65°C. Antibodies used for transcription factor ChIPs were rabbit anti-TTF1 (NKX2-1) (10 μg; Sigma-Millipore 07-601), and rabbit anti-PROX1 (10 μg; Proteintech 11067-2-AP).

Libraries were prepared using NEBNext® ChIP-Seq Library Prep Master Mix (E6240, NEB) and NEBNext® Multiplex Oligos for Illumina® (Index Primers Set 1 E7335S, Index Primers Set 2, E7500S) with Agencourt AMPure XP magnetic beads (Beckman coulter A63881) exactly as per manufacturer’s instructions. Libraries were checked using a Bioanalyzer and sequenced on a Hiseq 2500 sequencing platform.

### HiChIP

HiChIP assays were performed as in (33). Briefly, ∼10 million cells were crosslinked with 1% Formaldehyde and lysed with Hi-C lysis buffer (10 mM Tris-HCl pH 7.5, 10 mM NaCl, 0.2% NP40) supplemented with protease inhibitors. Chromatin was digested with the MboI restriction enzyme. The digested DNA sticky ends were filled with dCTP, dGTP, dTTP and biotin-labeled dATP by DNA Polymerase I, Large Klenow Fragment, and then ligated by T4 DNA ligase. Chromatin was sonicated in Nuclear Lysis buffer (50 mM Tris-HCl pH 7.5, 10 mM EDTA, 1% SDS) and then processed with regular chromatin immunoprecipitation procedures using the H3K27ac antibody (Abcam, ab4729). The ChIP-enriched Biotin-labeled DNA fragments were then purified by Streptavidin C1 magnetic beads and processed with Nextera DNA library preparation. The DNA libraries were sequenced by Illumina NextSeq (75 bp paired end reads).

### Bioinformatics analyses

#### Sequencing data analysis

ChIP-seq was demultiplexed using sample-specific Illumina primer sequences. The resulting data sets were aligned to the hg19 genome using Bowtie2 v2.2.6 (77). Reads with a Bowtie2 quality score less than 10 were removed using SAMtools v1.3 (78) with parameters (-bh -F 0×04 -q 10). Duplicate reads were removed using picardtools v1.119 (http://broadinstitute.github.io/picard), and the remaining reads were normalized to 10 million reads using HOMER v4.7 (47). All UCSC Genome Browser plots shown (79-82) reflect these normalized tag counts.

RNA-seq sample reads were aligned to the human genome using TopHat 2.1.0 (77, 83, 84). Default settings were used, with the exception of –G, specifying assembly to the hg19 genome, -- library-type fr -first strand, and –no-novel-juncs, which disregards noncanonical splice junctions when defining alignments. Cufflinks v2.2.1 (85-88) was used to call and quantify the transcripts from biological replicates for each sample, and identify genes, which were differentially expressed between samples, using the default parameters.

#### Peak calling

H3K27ac or TF ChIP-seq datasets from the SCLC samples were used to identify putative binding sites (ChIP-enriched peak regions) genome-wide based on the distribution of the aligned reads. ChIP-seq data sets for the transcription factors and the H3K27ac marks were normalized to 10 million reads. Peaks for each sample were called based on respective input samples created during immunoprecipitation. Peak calling was performed using the findPeaks library of HOMER 4.9 (47) and identified significantly enriched peak regions from each data set.

#### De novo motif discovery

HOMER v4.7 was used for all motif discovery presented here, To screen for potential DNA-binding co-factor motifs, a broader window of 150bp centered on the peak apex was used to identify enriched motifs adjacent to the primary binding site. The interval used for each analysis is noted in the text. Parameters –S 10 –bits were used unless otherwise noted. In de novo motif analysis, HOMER searches for motifs with 8, 10, or 12 bps on both strands such that flanking site preferences are not degraded.

#### SE identification

SEs were determined using H3K27Ac samples. H3K27ac bound enhancer regions that were significant at poisson p-value threshold of 1E-09 were identified using HOMER v4.7. Using ROSE (21, 26) tool enhancers within 12.5 kb were stitched together and rank ordered based on the ChIP signal versus input to identify SEs.

#### HiChIP data analysis

HiChIP reads were mapped to hg19 using HiC-pro (v.2.11.3) (89). For alignment, MboI restriction sites in the hg19 build were used. HiC-pro uses Bowtie2 for mapping and we specified --very-sensitive -L 30 --score-min L, -0.6, -0.2 --end-to-end –reorder for global options and --very-sensitive -L 20 -- score-min L, -0.6, -0.2 --end-to-end –reorder for local options. We used “GATCGATC” as the ligation site during the mapping process. We used H3K27ac enriched regions and hichipper (v.0.7.0) (90) to call loops. In this step we identified shared H3K27ac enriched regions in 6 out of 11 ASCL1 high samples and merged them to make an anchor regions set for hichipper to call loops from 8 ASCL1 high HiChIP samples. Similarly, we identified shared H3K27ac enriched regions in 4 out of 6 NEUROD1 high samples and merged them to make an anchor regions set for hichipper to call loops from 5 NEUROD1 high HiChIP samples. For Dual high HiChIP samples we identified shared anchor regions from the above ASCL1 high samples and NEUROD1 high samples and used this set as anchor regions to call loops in the 2 dual high HiChIP samples. From the hichipper results, we filtered the loops that are significant at a p-value of 0.01 and supported by at least 10 paired-end tags (PETS) for further analysis. For these filtered loops, we assigned a gene target to both loop ends if either of the loop coordinates fell within 5kb around the TSS of refseq genes. Similarly, if any anchor region fell within 5kb around the TSS of refseq genes we assign that gene to the respective anchor region.

#### Target calling for SE and TF ChIP peaks

The genes assigned to loop coordinates and anchor regions from the above step were used to identify SE-associated targets and putative targets for TF ChIP peaks. For this we compared the coordinates of SE regions or TF ChIP peaks with loop coordinates and anchor regions that are assigned with gene targets to identify the over lapped regions and used this information to assign gene targets for a given SE region or TF ChIP peak. Using this approach, we called the SE-associated targets for 11 ASCL1 high samples and 6 NEUROD1 high samples. For ASCL1, NKX2-1, and PROX1 ChIP peaks, we used a similar method to call the putative targets. Intersectbed module from bedtools (v2.27.1) (91) was used for assigning the gene targets to loop coordinates, and comparing them SE regions and TF peaks. For POU2F3 high sample SE regions, since there is no HiCHIP data available, we used gene targets that overlapped the SE regions or two nearest genes if SE regions are not overlapping with any gene targets.

#### Target comparison and filtering for pathway enrichment analysis

SE-associated genes from 11 ASCL1 high samples were compared and those shared between 6 of 11 ASCL1 high samples were identified as SCLC-A SE-associated genes. A similar strategy was used for SCLC-N (4 of 6) and SCLC-P (3 of 5). To focus on abundantly expressed SE-associated genes, we used an expression cutoff of 5 FPKM in a majority of the relevant SCLC subtype. These filtered data sets used for pathway enrichment analysis.

ASCL1, NKX2-1 and PROX1 ChIP peak associated targets in NCI-H2107, NCI-H889, and NCI-H128 cells were compared and putative targets identified if they were shared in 2 out of 3 cell lines for each factor. These gene lists were compared with ASCL1-KD and 3TF-KD RNA-seq DEG lists and pathways enrichment analysis performed. All pathway enrichment analyses in this study were performed using gprofiler tool (92) using GO biological processes, reactome, KEGG and wiki pathway databases.

For the GSEA enrichment map in Fig. 4E, a gene set enrichment analysis (GSEA) was performed (Subramanian et al., 2005) from ranked list of DEG upon ASCL1 and the 3TF knock down in NCI-H2107 cells. Default parameters were used for GSEA except the permutation type parameter. Due to the limited number of samples, we used the “geneset” permutation type for the analysis. We performed GSEA for each condition separately and the gene sets that were significantly enriched in (FDR ≤ 5%) at any given condition were selected for clustering analysis. Hierarchical clustering was performed on the normalized enrichment scores (NES) obtained from the significant gene sets enriched in the two conditions and plotted as an enrichment heatmap using heatmap.2 function available in “gplots” R package (93).

### Immunoprecipitation

Nuclear fractions from ∼1×10^6^ NCI-H2107 cells were prepared using the NE-PER kit (ThermoFisher Scientific 78835) as per the manufacturer’s instructions and incubated with 10 μg primary antibodies overnight at 4°C followed by 90 minutes incubation with 35 μl protein A/G agarose plus (50% slurry) (ThermoFisher Scientific 20421). Immune pellets were washed 5 times in PBS containing 1% Triton-X100 and analyzed by SDS-PAGE followed by either immunoblotting or mass-spectrometry (LC/MS-MS). For LC/MS-MS, 6 immunoprecipitations were pooled. Antibodies used for immunoprecipitations were mouse anti-ASCL1 antibodies (BD-Biosciences 556604) and normal mouse IgG (Santa Cruz Biotechnology sc-2025). For Figure 3B, a guinea pig anti-ASCL1 antibody was used (TX518, (94)). Nuclear extracts of NCI-H524 cells, which do not express ASCL1, PROX1, or NKX2-1, were used as negative controls.

### Mass-spectrometry analysis

For protein identification, samples were separated by SDS-PAGE electrophoresis. After Coomassie stain, bands were cut out of the gel and submitted to the UT Southwestern Proteomics core facility for LC-MS/MS analysis on an Orbitrap Fusion Lumos mass spectrometer. Proteins were identified using Proteome Discoverer 2.2.

### Immunoblotting

Cellular extracts prepared as in (95) and immune pellets were analyzed by SDS-PAGE electrophoresis using BIORAD Criterion gels and electro-transferred on PVDF membrane (EMD-Millipore IPVH85R). Membranes were incubated overnight at 4°C with primary antibodies and for 2 hours at room temperature with horseradish-conjugated secondary antibodies. Western-blots were detected using the SuperSignal™ West Pico PLUS Chemiluminescent Substrate (ThermoFisher Scientific 34580). Western-blots were quantified using Image Studio Lite quantification software (Licor).

Antibodies used for immunoblotting include mouse anti-ASCL1 (1:1000; BD-Biosciences 556604), rabbit anti-NKX2-1 (1:2000; Cell Signaling Technology 12373), rabbit anti-PROX1 (1:2000; Proteintech 11067-2-AP), rabbit anti-NFIB (1:4000; Abcam Ab186738), rabbit anti-FOXA2 (1:1000, Cell Signaling Technology 3143), rabbit anti-SCN3A (1:500; Santa Cruz sc-517010), rabbit anti-KCNB2 (1:500; Santa Cruz sc-376275), mouse anti-GAPDH (1:10000, Santa Cruz Biotechnology sc-47724) and rabbit anti-β-tubulin (1:2000, Santa Cruz sc-5274). Secondary antibodies were HRP-conjugated goat anti-Rabbit IgG (H+L) (1:10000, ThermoFisher Scientific 31460) and HRP-conjugated goat anti-Mouse IgG (H+L) (1:10000, ThermoFisher Scientific A16066).

### Cell surface staining and flow cytometry

NCI-H2107, NCI-H524, and NCI-H526 cells (∼2×10^6^) were harvested by centrifugation and resuspended in ice-cold FACs buffer (PBS, 10% FBS, 0.1% Sodium Azide). SCLC mixed samples contained ∼10^6^ cells of each cell line mixed immediately prior to staining. Cells were incubated with SCN3A antibody (Alomone Labs, Anti-SCN3A (Nav1.3) (extracellular), Cat #: ASC-023, 4.25 µg/ml) for 30 min on ice, washed 3 times with ice cold FACs buffer, incubated with Alexa Fluor 568-conjugated secondary antibody (Thermofisher, Catalog # A10042, 1:400) for 30 min on ice, washed 3 times as above, and resuspended in ice cold FACs buffer. A FACS Aria II SORP machine using a 561 nm laser separated positive from negative cells. Cells incubated with secondary antibody only were used to control for background fluorescence. Raw data was analyzed using FlowJo 10.7.0.

### RT-qPCR

RNA was extracted from cell pellets, or Alexa Fluor 568+ and – cell fractions sorted into Trizol®, using the Arcturus PicoPure RNA kit as described above. cDNA was synthesized using the SuperScriptTM III First-Strand Synthesis SuperMix (Invitrogen,™ Cat. # 11752050). RT-qPCR was performed on an Applied Biosystems® 7500 Fast thermal cycler using Fast SYBRTM Green Master Mix (Applied Biosystems®, Cat. # 4385612). All reactions were performed in triplicate. Primers used: *SCN3A* Forward (5’ GGAGAGCTGTTGGAAAGTTCTT-3’), *SCN3A* Reverse (5’-TTCCTTCGGTTCCTCCATTCT-3’), *ASCL1* Forward (5’-CCCAAGCAAGTCAAGCGACA-3’), *ASCL1* Reverse (5’-AAGCCGCTGAAGTTGAGCC-3’).

### Soft agar colony formation assay

NCI-H889, NCI-H524, and NCI-H526 cells were seeded at 20,000 cells/well in 12-well plates in a layer of 0.32% agar/RPMI/FBS/0.64% Antibiotic-Antimycotic (Thermofisher) over a bottom layer containing 0.5% agar/RPMI/FBS/0.64% Antibiotic-Antimycotic. Cells were treated with vehicle, ICA-121431 (Sigma #SML-1035), or Quinine hydrochloride (Tocris #4114) at 1-100µM in the top agar layer at the time of plating. Cultures were maintained for 2.5 weeks in a humidified, 37°C incubator containing 5% CO2 and 95% air. Colonies were then fixed and stained with 14.3% ethanol and 0.1% crystal violet in water. Plates were imaged by light microscopy using a Keyence BZ-X700 Series microscope, and colonies quantified using ImageJ.

### Statistics

Quantification of data obtained from WST-1 assays and immunoblots is reported as mean ± SEM and the sample size is as indicated. For Fig. 4B’-C and Fig. 6F, P values were determined using one-way ANOVA followed by a Bonferroni post-hoc test. For Fig. 3C and Fig. 6A, the non-parametric test, Mann-Whitney, was used to determine P values. Graphpad Prism v7.0 was used for statistical analysis.

### Accession numbers

The accession number for the RNA-seq, ChIP-seq and HiChIP data generated for this project is GSE151002. Other data sets used include H3K27ac: GSE85401 (HBE), GSE72956 (NCI-H2087), GSE89128 (A549, NCI-H3122, PC9), GSE115124 (DMS79, NCI-H146, NCI-H128, NCI-H446, CORL311, DMS114), GSE62412 (GLC16, H82 Input). Also, for ASCL1-ChIP: GSE69394, and for RNA-seq from the SCLC cell lines: dbGaP Study Accession phs001823.v1.p1

## AUTHOR CONTRIBUTIONS

Conceptualization, K.P. and J.E.J.; Investigation, K.P., K.R., D.P.K. and X.Z.; Data analysis, R.K.K.; Resources, J.E.J., J.D.M., and X.Z.; Writing, K.P., R.K.K. and J.E.J.; Funding Acquisition, K.P., J.E.J. X.Z. and J.D.M; Supervision, J.E.J.

## ACKNOWLEDGEMENTS

We acknowledge the many helpful discussions with members of the SCLC interest group at UT Southwestern, Chaoying Liang in the Microarray Core at UT Southwestern for Next Generation Sequencing and Mohammad Goodarzi in the Proteomics Core facility at UT Southwestern for outstanding service. We thank Victor Stastny, Kenneth Huffman, Luc Girard, and Adi Gazdar (deceased) for work on the SCLC lines and discussions about SCLC lineage transcription factors, and Charles Rudin and J.T. Poirier for providing the SCLC PDX. Support for maintaining, authentication, and genomic characterization of the SCLC cell lines was provided in part by NCI U24CA213274. Funding for this project was provided by the NCI including U01CA213338 to J.D.M., R00 CA215244 to X.Z, a Career Development Award from the Spore Grant in Lung Cancer P50CA70907 to K.P., F30 CA228314 to D.P.K, and the Cancer Prevention Research Institute of Texas Training Grant RP160157 to K.R.

